# Transcriptome analysis of potato leaves in response to infection with the necrotrophic pathogen *Alternaria solani*

**DOI:** 10.1101/2020.09.21.307314

**Authors:** Mengjun Tian, Jinhui Wang, Zheng Li, Chen Wang, Dai Zhang, Yiqing Yang, Yang Pan, Dongmei Zhao, Zhihui Yang, Jiehua Zhu

## Abstract

The potato early blight was caused by *Alternaria solani* (Aso). At present, potato early blight resistant varieties are lacking. The experiment of Aso inoculation on a.k.a. Favorita showed that the content of chlorophyll decreased continuously within 2 days after inoculation. In addition, the contents of JA-Ile, SA, ABA and IAA were determined by LC-MS.The results of two defense-related hormones showed that jasmonic acid content decreased significantly, while jasmonic acid content did not change significantly. In order to understand the difference in gene expression of potatoes infected by Aso, the 24 hpi, 36 hpi and 48 hpi transcriptome were determined. The results showed that the gene annotation and cluster analysis of DEGs revealed a variety of defense responses to Aso infection, especially the plant hormone signal transduction pathway. The differential expression of 10 DEGs was confirmed by qPCR analysis. Finally, through weighted gene co-expression network analysis (WGCNA), “samonl” was determined as a key hormone-related network in the process of Aso infection in potatoes. This study provides an important clue for understanding the interaction between potato and Aso and for disease resistance breeding.

## INTRODUCTION

The early blight (EB) of potato caused by *Alternaria solani* (Aso) is one of the most serious fungal leaf diseases of potato [1]. After the potato leaves infected by Aso, usually results in the formation of leaf necrosis, and caused chlorotic around the lesion [2]. With the photosynthetic potential to reduce disease development and lead to leaf premature senescence and advance into death [2]. The relatively slow destruction of host tissues has a serious impact on the normal growth and development of crops and yields [3]. According to the research, the annual production loss caused by early blight damage is up to 50% [4].

In most crops, plant photosynthesis can effectively accumulate carbohydrates (such as sugar and starch), and to promote the growth and development of fruits and seeds [5]. With the deepening of research, the change of chloroplast activity is crucial for the survival or elimination of plant pathogens. For example, the infection of Brassica juncea by the fungal pathogen *Alternaria brassicicola* severely disturbs chloroplast morphology and function. [6]. It seems that the survival of pathogens does not only cause structural damage to chloroplast. During the onset of disease, chloroplasts resist pathogens by producing immunogenic signals in the form of ROS, Ca^2+^ and plant hormones [7,8]. For pathogens, they can successfully survive by eliminating the plant’s immune program and promoting signals that are beneficial to themselves [9]. Plant hormone mediated signaling pathways play an important role in plant disease resistance [10], when plants detect the invasion of pathogens, they quickly start the complex hormone signal network for pathogen resistance [11]. The life style of pathogens makes different plant hormone signaling pathways present a variety of responses [10]. In general, JA signaling pathway enhance the resistance to hemi-biotrophic and necrotrophic pathogens, while the resistance to biotrophic pathogens mainly depends on SA mediated signaling pathways [12,13]. Previous studies have suggested that SA and JA signal transduction pathways have antagonistic effects [14]. Interestingly, recent studies have shown that SA or JA signaling pathways do not independently mediate resistance to biotrophic or necrotrophic pathogens. At present, in several researches on necrotrophic pathogens, SA and JA signaling pathways are involved in the defense response of plants against necrotrophic pathogens [15,16,17]. In order to understand the mechanism of pathogen invasion, transcriptome is an effective strategy to study the disease system. At present, some transcriptome studies on host invasion of *Alternaria sp*. have been reported. In one study, the hormone content of chrysanthemum inoculated with *Alternaria sp*. showed that JA and SA contents increased, and defense enzyme activities in infected plants increased significantly. Transcriptome sequencing of leaves infected with *Alternaria sp*. for 3 and 5 days showed that JA and SA signaling pathways were highly responsive [17]. In another study, used disease bioassays and transcriptomic analysis showed that intact SA-signalling is required for potato defences against the necrotrophic fungal pathogen *Alternaria solani* [18]. Although several studies have been carried out to analyze the interaction between *Alternaria solani* and host plants [17,18,19], this study is the first global transcriptome analysis of potato infected by *Alternaria solani*. In this study, we tried to understand the change of transcriptional expression related to potato defense necrotrophic pathogen by bioassay of diseases in potato leaves, content determination of several hormones such as jasmonate and salicylic acid, and transcriptome analysis of infected potato leaves the transcriptional response of delicious to Aso infection and the potential cause of its susceptibility.

## Results

### Phenotypic effects of *Alternaria.solani* infection in potato leaves

Potato plants (cultivar Dutch 15, a.k.a. Favorita) were inoculated in greenhouse with Aso, by foliar sprays with spore suspensions of Aso isolate HWC-168. Symptoms of early blight appeared on the inoculated leaves at 24 hours post-inoculation (hpi), and disease symptoms developed rapidly in the subsequent 24 hours (Figure 1; Supplementary Figure S1). Mild chlorosis was observed at 24 hpi, which turned into general chlorosis with dark necrotic spots at later time points (Figure 1). At 4 days post inoculation, potato leaves inoculated with Aso has turned completely yellow with necrotic patches (Supplementary Figure S2), indicating the leaves had undergone Aso-induced premature senescence.

**Figure 1.**
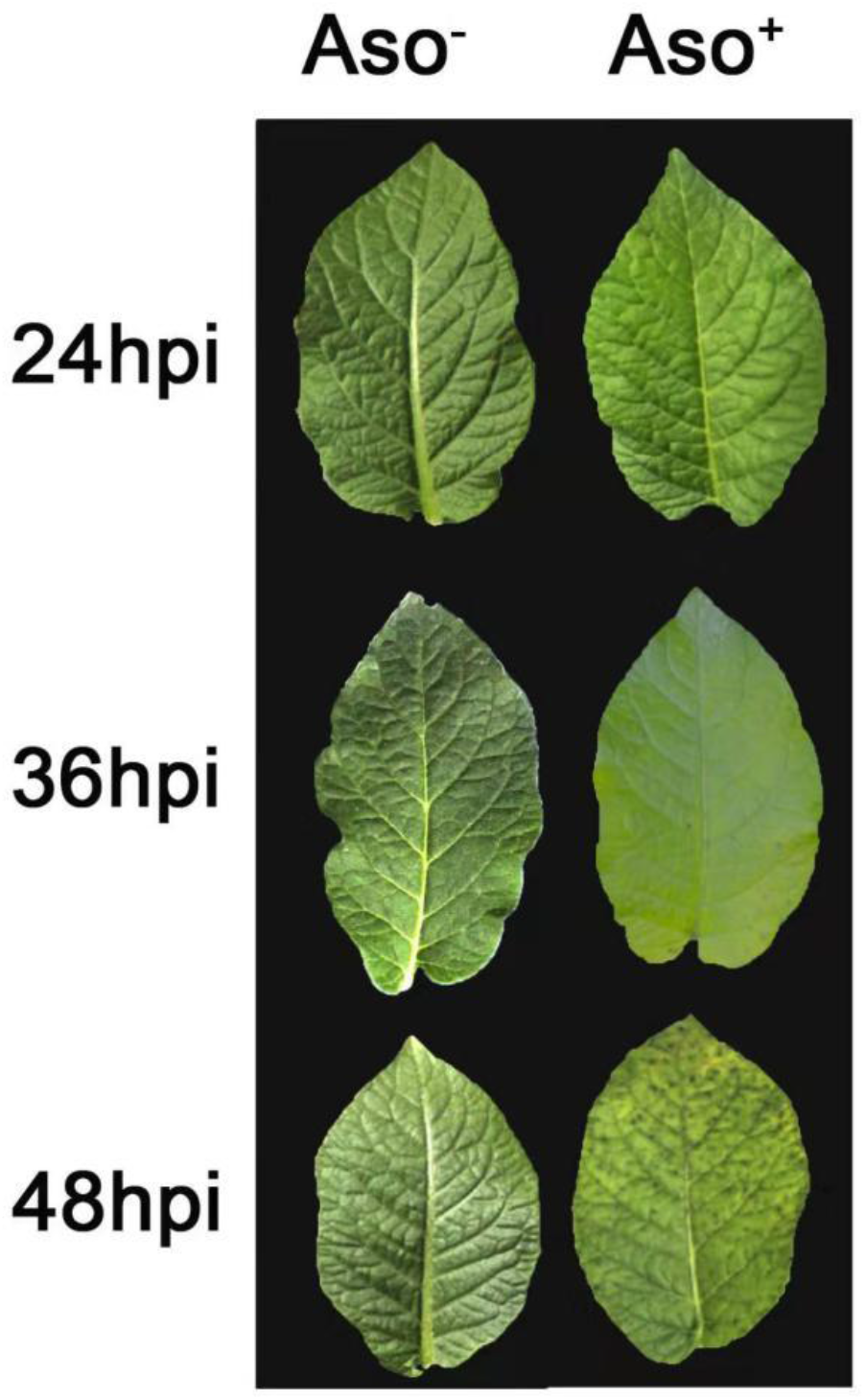
Changes in symptoms of early blight in potato leaves after inoculation with *Alternaria solani* (Aso) isolate HWC-168. The photos were taken at 24, 36 and 48 hours post-inoculation. Leaves on the left (Aso^-^) were inoculated with 0.1% Tween 80 (mock); leaves on the right (Aso^+^) were inoculated with spore suspension of Aso isolate HWC-168.

### Altered chlorophyll and plant hormone levels in the infected leaf tissues

Aso infection caused a reduction in the content of total chlorophylls (chlorophyll a and b) in potato leaf tissues. Although no significant difference (p-value 0.229) was found in the chlorophyll content between the infected (1.488±0.012 μg/gFW) and control (1.532±0.017 μg/gFW) plants at 24 hpi (Figure 2). However, more drastic reductions in total chlororphyll content were detected at later time points. The chlorophyll contents in the Aso-infected leaves at 36 hpi and 48 hpi were 1.144±0.051 μg/gFW and 0.672±0.065 μg/gFW, respectively (Figure 2). This result coincides with phenotypic alterations in the infected leaves that typical chlorosis symptom was visible at later time points (Figure 1).

**Figure 2.**
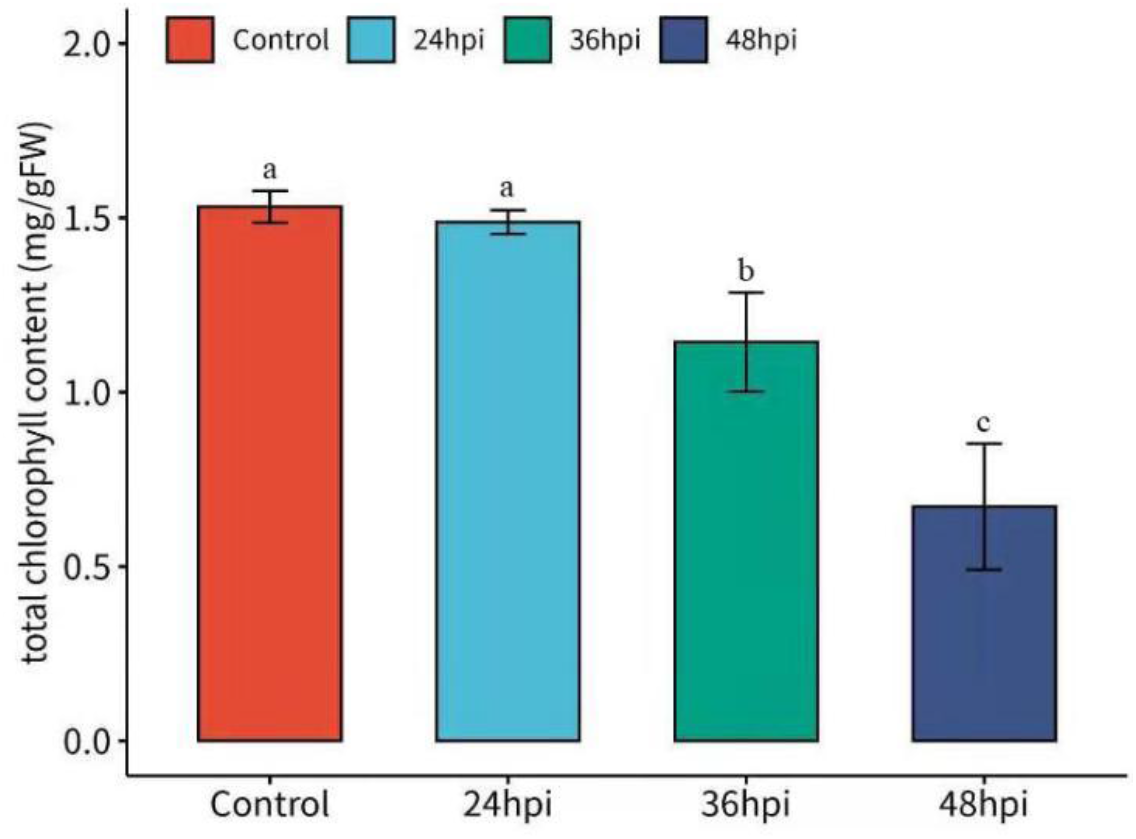
Effects of *Alternaria solani* (Aso) on the total chlorophylls content of potato leaves. The chlorophylls contents were significantly decreased at 36 and 48 hours post-inoculation in the Aso-infected plants. Bars with different letters indicate significant differences between groups (*p* <0.01, Welch’s ANOVA followed by Games-Howell post-hoc test), and error bars indicate 95% confidence interval. FW, fresh weight.

The concentrations of salicylic acid (SA), abisisic acid (ABA), indole acetic acid (IAA), and jasmonoyl isoleucine (JA-Ile) in Aso-infected and non-infected leaves were measured. The concentration of ABA in the infected leaves were significantly lower than in the control plants at all time points (Figure 3a). The concentration of IAA was significantly lower in the infected leaves than in the non-infected leaves at first two time points but not at 48 hpi (Figure 3b). The concentration of JA-Ile in the infected leaves was lower than in the control plants at 36 and 48 hpi but not at the first time point (Figure 3c). No significant changes in SA concentration was detected between the infected and control plants at any time point (Figure 3d).

**Figure 3.**
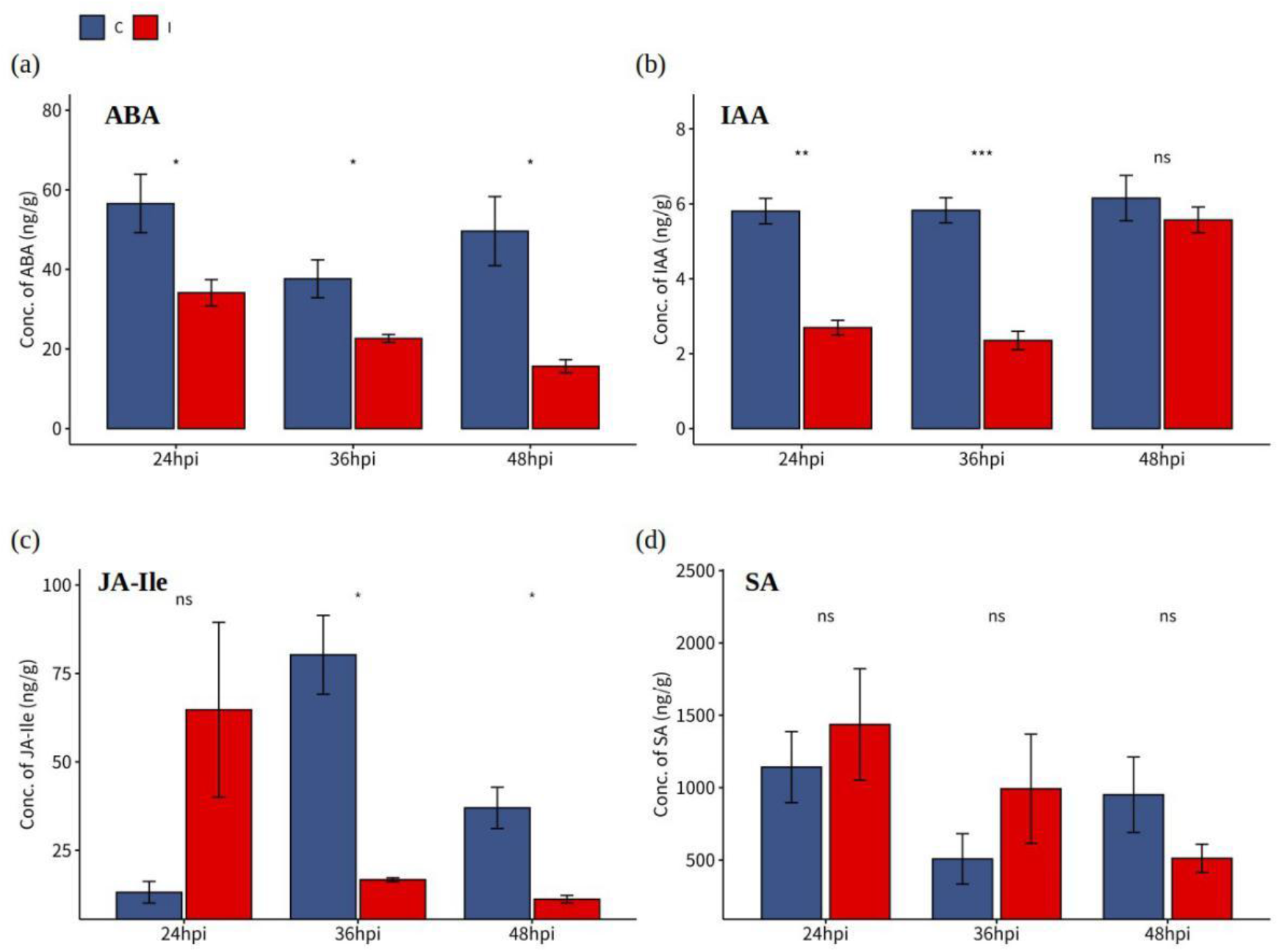
Plant hormone levels in potato leaves of mock inoculation (C) and *Alternaria solani* (Aso) infected (I) plants at 24, 36 and 48 hours post-inoculation. ABA, abscisic acid; IAA, indole acetic acid; JA, jasmonic acid; SA, salicylic acid. Asterisks indicate statistical significance determined by Student’s t-test at \**p* <0.05, \*\**p* <0.01 and \*\*\**p* <0.001; ns represents no significance (*p* >0.05). Bars represent standard errors.

### RNA-Seq datasets and differentially expressed genes in the infected potato plants

On average, each RNA-Seq sample yielded 24.73M read pairs, and the clean dataset had an average of 75.24% overall mapping rate (Supplementary Table S1). The result of principal component analysis (PCA) of the RNA-Seq datasets showed that the first principal component (PC1) and PC2 explained 76% and 11% of the total variation, respectively (Supplementary Figure S3). The PCA plot showed that the infected samples and the control samples clustered into different groups which agree with the experimental design. In this experiment, 6184, 10887 and 8109 differential expression genes (DEGs) of potato were identified by comparison between the infected and control plants at the three post-inoculation time points, respectively (Supplementary Table S2). More DEGs were identified at 36 hpi suggesting other environmental factors (mainly photoperiod) also affect gene expression during disease development (Supplementary Figure S3).

### DEGs involved in leaf pigment biosynthesis and photosynthesis

Aso infection caused expression levels of many genes involved in leaf pigment synthesis and photosynthesis were down-regulated at all time points. DEGs related to chlorophyll biosynthesis and photosynthesis were enriched in predefined gene sets, including Gene Ontology (GO) terms “photosynthesis”, “thylakoid”, “generation of precursor metabolites and energy”, “cellular protein modification process” and Kyoto Encyclopedia of Genes and Genomes (KEGG) pathways “photosynthesis”, “antenna proteins”, “photosynthesis proteins”, “carbon fixation in photosynthetic organisms” (Supplementary Table S3). In general, more drastic down-regulations of those genes were detected at 36 hpi than at other two time points (Figure 4), which suggests that the function of chloroplast was strongly suppressed by Aso infection at day time (36 hpi).

**Figure 4.**
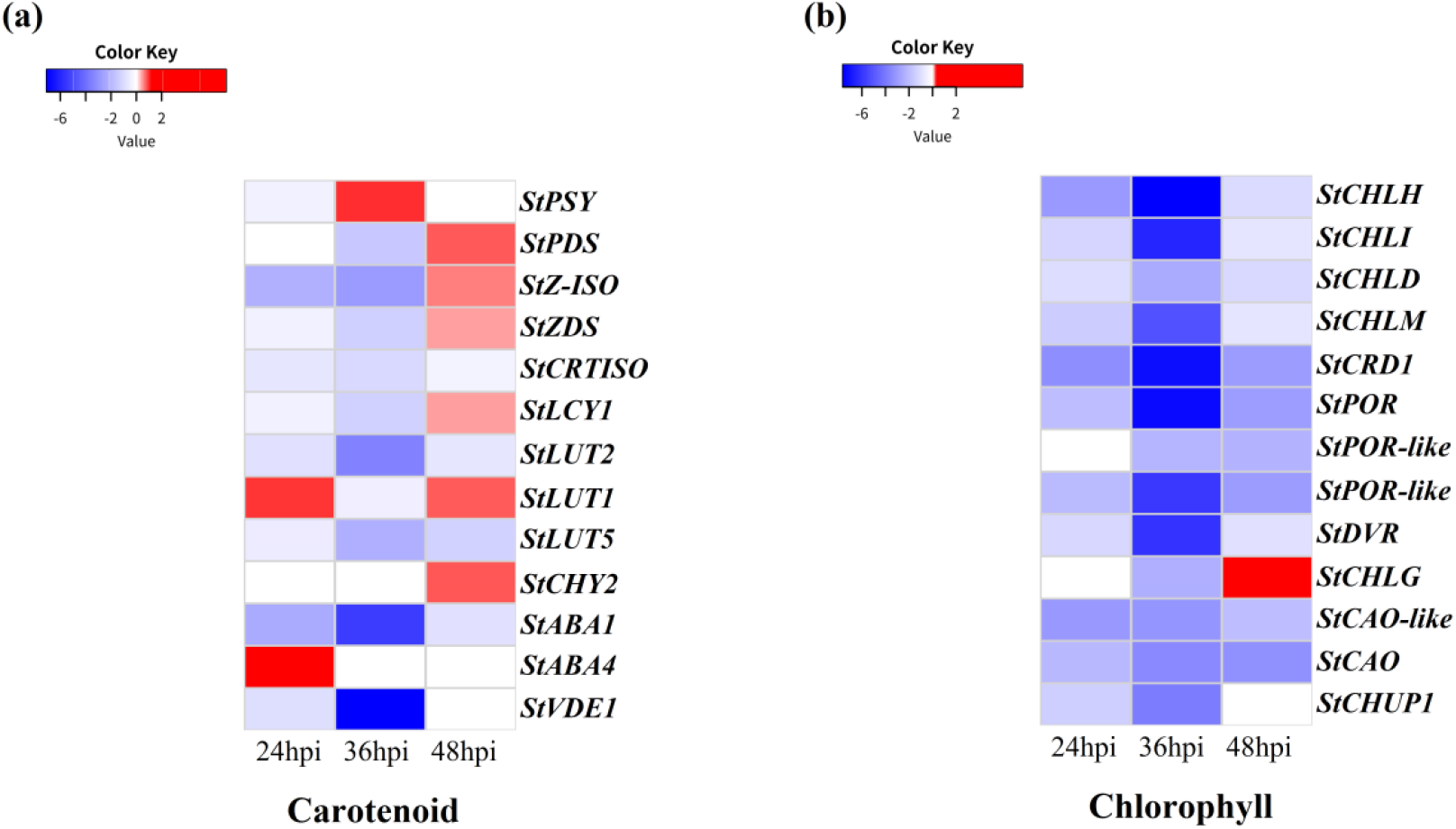
Heatmaps of DEGs involved in leaf pigment biosynthesis pathways, including chlorophyll biosynthesis and carotenoid biosynthesis pathways. The log2|Foldchange| was colored using Cluster 3.0 (red for up-regulated, blue for down-regulated), each horizontal row represents a DEG with its gene name, and the vertical columns represent 24, 36, and 48 hpi from left to right.

Most of the genes encoding enzymes involved in chlorophyll biosynthesis were identified to be down-regulated in infected potato leaves (Figure 4b), and the down-regulation was more intense at 36 hpi, which was consistent with the significant decline of total chlorophyll content measured at 36 hpi (Figure 2). However, the expression levels of carotenoid and lutein biosynthesis related genes were different at three time points after inoculation (Figure 4a). Most genes related to carotenoid biosynthesis were down-regulated at 36 hpi, but up-regulated slightly at 48 hpi. The effect of Aso infection on lutein synthesis at 24 and 48 hpi is difficult to predict because *LUT1, LUT2* and *LUT5* show inconsistent expression patterns across time points. Nevertheless, *LUT1* and *LUT2*, the main regulators of lutein biosynthesis pathway, were all significantly down-regulated at 36 hpi, therefore lutein biosynthesis was suppressed at 36 hpi. In addition, most genes encoding antenna complex and photosystem subunits were down-regulated after Aso infection (Supplementary Table S4), suggesting the Aso infection has greatly hampered the photosynthesis in potato plants at very early time point.

### DEGs involved in autophagy and leaf senescence

In Arabidopsis, *WRKY53* and *NAP2* are two transcription factors (TFs) as markers that are tightly associated with senescence [20]. The potato homolog of *AtNAP2* was up-regulated at all time points, and homologs of *AtWRKY53* were up-regulated at 24 and 48 hpi but not at 36 hpi. Some enzymes involved in chlorophyll breakdown and degradation during leaf senescence, such as *SGR1, PPH, PAO, RCCR* were up-regulated (Supplementary Table S4). A large number of up-regulated expressions of leaf senescence-related genes serve as transcriptional evidence to explain the strong early leaf senescence observed 4 days after Aso infection (Supplementary Figure S2). Most autophagy related genes were significantly up-regulated at 36 hpi, including genes involved in initiation of autophagy (*ATG1* and *ATG13*), formation of autophagosome and fusion with the vacuole (*ATG6, VPS15, VPS34, VTI12*), and also genes that are components of the *ATG9* complex (*ATG2, ATG9, ATG18*) (Supplementary Table S4). But an exception is the initial regulator (*TOR*) [21], its expression levels were not altered by Aso infection at all time points.

### DEGs involved in plant hormone biosynthesis and signal transduction

Aso infection significantly altered the expression levels of genes involved in plant hormone biosynthesis and signaling pathways that were enriched in gene sets including GO terms “DNA-binding transcription factor activity”, and “transcription regulator activity”, “response to endogenous stimulus”, and KEGG pathways “MAPK signaling pathway”, “phenylalanine metabolism”, “alpha-linolenic acid metabolism”, “cysteine and methionine metabolism” and “plant hormone signal transduction” (Supplementary Table S3).

The expressions of genes involved in ABA biosynthesis, *ZEP* (*ABA1*), *ABA2*, and *NCED1*, a rate limited enzyme in Arabidopsis [22], were down-regulated at three time points (Supplementary table S4), which partly explained the reduced ABA content in the infected leaf tissue (Figure 3a). Most *PYL* receptors in ABA signaling pathway were up-regulated in infected leaves, while most *PP2Cs*, the negative regulators of ABA signaling, was down-regulated in infected plants. The expression of *JA2*, a transcription factor involved in ABA-mediated stomatal closure, was constantly up-regulated at three time points. However, as a signal activator of ABA, *SNRK2* has not been found common differential expression pattern across time points.

The expression of *ACO* and *ACS* in Aso infected leaves increased significantly at 36 hpi and 24 hpi, respectively. In general, the genes in ETH signal transduction pathway were up-regulated but not down-regulated. Two out of three *EIN3/EIL* were up-regulated at 24 and 36 hpi, *EBF1/2* was up-regulated at 24 and 48 hpi, and *EIN5* was up-regulated at 36 hpi (Supplementary Table S4).

In Arabidopsis, during pathogen infection, salicylic acid is mainly synthesized through chorismate ICS1 (isochorisate synthase I) pathway in the chloroplast [23]. Data analysis showed that the expression of *ICS1* was continuously suppressed at three time points. However, *PAL* and *PAL-like* genes were up-regulated in infected leaves (Figure 5a). In the process of salicylic acid biosynthesis, *EDS5* is responsible for transporting SA precursor substances from the chloroplast plastid to the cytoplasm and is catalyzed by PBS3 to generate isochorismate-9-glutamate and further spontaneously decompose to generate SA [24]. The homologous gene of *PBS3* has not been cloned in potato, but in our data we found that *EDS5* showed down-regulated at 36hpi. *PR1*, as the maker gene of SA-mediated Pathogens resistance, is regulated by NPR1 [25]. Although the actual measurement data of SA content did not show significant differences at the three time points after inoculation with Aso, the downstream genes of SA signal transduction (*NPR1, TGA2.1, PR1, PR5*) were up-regulated in the three times after Aso inoculation.

**Figure 5.**
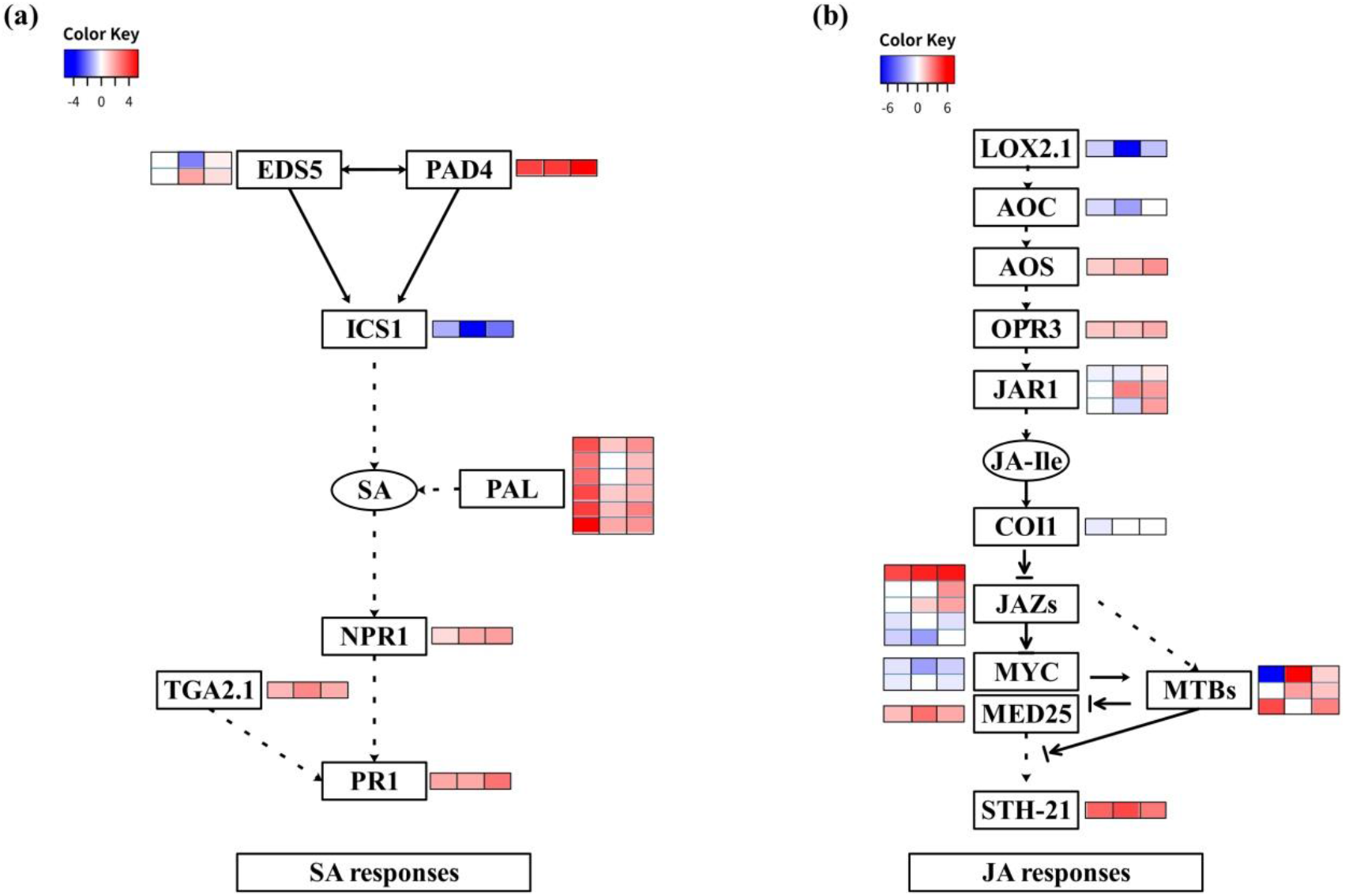
Gene expression model of SA and JA response pathway after Aso inoculation. (a) SA response pathway model. (b) JA response pathway model. The heat map shows the changes in gene expression at 24h, 36h, and 48h after Aso inoculation.

Although the genes encoding *AOS1* and *OPR3*, which involved in the JA biosynthetic pathway were up-regulated, but the two key rate-limited enzyme genes, *LOX2.1* and *AOC* [26] (Figure 5b) were significantly down-regulated. Combined with the JA-ILE measurement data (Figure 3c), it is speculated that the JA biosynthetic pathway was suppressed at three time points. The master regulater of JA Signal transduction pathway is *MYC2* [27]. *MYC2* occupies a high level in the JA signal transduction pathway and can actively induce the transcription of defense genes [27]. Although the *stMYC2* were not differentially expressed, *stMYC1* was significantly suppressed at three time points. *MTBs* plays the role of loop regulated *MYC2* in JA signal transduction [28]. The physical interaction between MYC2 and the MED25 subunit of the mediator transcription co-activator complex determines the function of *MYC2* [29]. Although *MYC2* is not differentially expressed, we found that *MED25* continues to be up-regulated at three time points. In our data, we found that the transcriptional expression patterns of *MTB1* and *MTB2* at the three time points of the three members of the *MTBs* family showed consistency, but the other member *MTB3* Showed the opposite mode of expression (Figure 5b). As a repressor of *MYC2*, we observed that except for *JAZ7* which was specifically down-regulated at 36hpi, the rest of *JAZs* (*JAZ2, JAZ3, JAZ5*) were up-regulated to varying degrees at three time points (Figure 5b).

### Validation of gene expression by qRT-PCR analysis

The expression of 10 DEGs was verified in qRT-PCR analysis to verify the accuracy and reproducibility of the transcriptome data. The DEGs for qRT-PCR verification represent different physiological aspects of potato plant, including plant hormone biosynthesis and signal transduction, pigment biosynthesis and defense responses, etc. In S A biosynthesis and signal transduction pathway, it includes *ICS1* and *NPR1*. In JA biosynthesis and signal transduction pathway selcect *LOX2.1, MYC1, MTB1* and *JAZ2*. Among the pigment biosynthesis pathways, we chose *POR1, POR3* and *ZEP* to verified. The expression level of all selected genes changed significantly after Aso inoculation. The expression patterns of these genes determined by qRT-PCR were similar to those obtained based on RNA-seq (Figure6).

**Figure 6.**
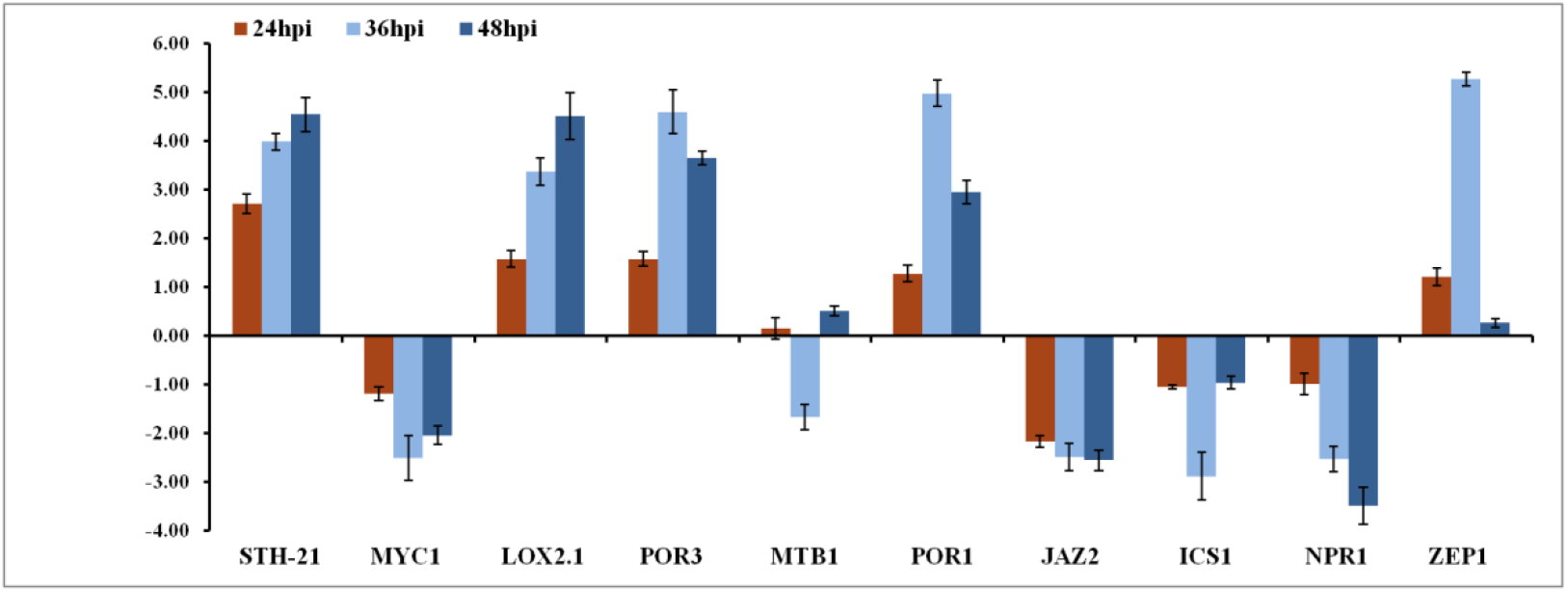
Using qRT-PCR to verify the transcription changes of 10 genes selected from DEG. These genes expression level were all determined by Real-time PCR. Vertical bars indicate the standard deviation of three biological replicates.

**Figure 7.**
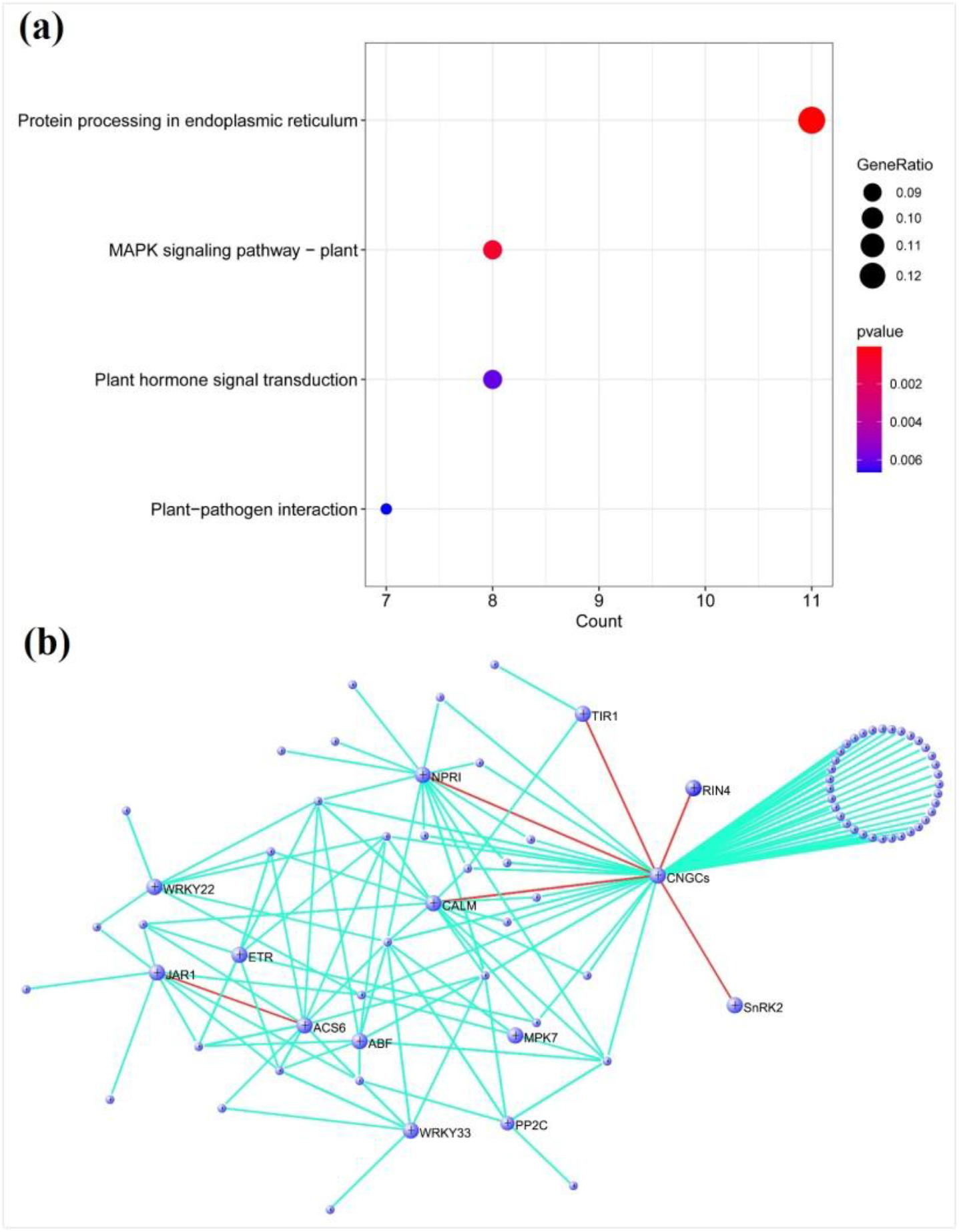
Network co-expression analysis of RNA-seq data. (a) KEGG analysis of “samonl” modular gene related to phytohormone phenotype. (b) Plant hormone-related phenotype “samonl” module, the enlarged circle represents the hub gene encoded according to its biological function, and the correlation between the hub genes is indicated by the red connecting line.

### Co-expression network revealed the reaction of phytohormone to Aso infection in potato

After removing genes with low expression and little overall change, a total of 6323 genes were obtained for WGCNA analysis. A total of 14 modules were generated in the analysis results, among which the “salmon” module was significantly related to plant hormones. Further KEGG analysis of the “salmon” module found that a total of 253 genes were enriched into four types of gene terms, include protein processing in endoplasmic reticulum, MAPK signaling pathway-plant, plant hormone signal transduction a nd plant-pathogen interaction. Through the analysis of the “samonl” module genes, w e can determine the plant-pathogen interactions (*WRKY33, CNGC, RIN4, ACS6 and C ALM*) involved, plant hormone signal transduction genes and seven genes (*PP2C, Sn RK2, ERS, TIR1, JAR1, ABF* and *NPR1*) appeared as hub genes in the network. In particular, the cyclic nucleotide gated channel *CNGCs* is associated with a large number o f genes in the module, as a channel protein that controls Ca2^+^, this gene is extremely important in the process of participating in plant immune responses [30]. Based on the above results, we determine that the “samonl” module is closely related to the process of plant hormone signal transduction and the interaction between plants and pathogens.

## Discussion

At present, there is a lack of comprehensive understanding of the pathogenic mechanism of potato early blight. However, with the release of the potato genome and the use of transcriptome sequencing technology, it is possible to fully understand this sensitive process at the transcriptome level. After inoculated potato plants in the greenhouse, we found that the potato leaves quickly turned yellow and entered the stage of senescence and death after Aso infection. Although several studies have reported on the pathogenic mechanism of Aso [18,19], we hope that through a comprehensive analysis of the host’s transcriptome response during the process of infection (from successful colonization to leaf necrosis), we can gain insight into the pathophysiology of the disease and other aspects.

The integration of rich KEGG pathways and GO analysis results enables people to have a deeper understanding of the different responses of different species and tissues to pathogen stress. Photosynthesis converts light energy into chemical energy in chloroplasts, which is very important for the normal growth and development of crops [31]. In our study, we found that after Aso infection, decreased chlorophyll content in potato leaves, potato leaves color has undergone significant changes (Figure 2). At the three time points after Aso infection, almost all GO terms “ generation of precursor metabolites and energy “ and “photosynthetic antenna protein” of the KEGG pathway were down-regulated (Supplementary Table S3). According to these processes, the three richest in GO terms are “optical system”, “chloroplast stroma” and “thylakoids” (Supplementary Table S3). Almost all genes are most strongly suppressed at 36hpi, and therefore susceptibility process can have a strong interaction with light that environmental factors during the day.

Through the complex network of plant hormone signaling pathways, plants can effectively respond to pathogens and balance the relationship between defense and growth [32]. To characterize the antagonism and co-operation of signal transduction among salicylic acid, jasmonic acid and abscisic acid can better develop economically important disease management strategies for crops [33]. Salicylic acid has been proved to have two important branches in the process of biosynthesis, including ICS pathway and PAL pathway [34]. We found that there was no significant change of SA content in Aso infected leaves(Figure3d). Interestingly, when we investigated the key genes in SA biosynthesis pathway in transcriptome data, we found that the key genes in the two branches showed opposite transcriptional expression patterns (Supplementary Table S2). Therefore, the compensation effect of the two pathways of SA biosynthesis may be the reason for the unchanged SA content. Salicylic acid can trigger the activation of downstream disease-related genes (PR) and provide a marker for the basic and induced defense responses of PTI and ETI [35]. In the transcriptome data, after infection with Aso, the genes related to pathogen infection (such as *NPR1, PR-1, STH-21*) downstream of SA are significantly activated (Supplementary Table S2). A recent study used microarray to characterize potatoes in order to cope with Aso infection, a complete SA signal transduction pathway is necessary [18]. Although our data shows that SA content did not change significantly after Aso infection, but its downstream defense response was significantly activated, so our study provide a more complete evidence, enrich the SA pathway response mechanism after the potato in response to infection by Aso.

In response to the invasion of necrotic pathogens, jasmonic acid is (+)-7-isojasmone-1-isoleucine (JA-Ile), which can be sensed through the COI1-JAZ co-receptor system in the disease-resistant response [36]. Although we did not observe that JA-Ile was inhibited at 24hpi, it was significantly inhibited at 36hpi and 48hpi. The JA signaling pathway plays an important role in resisting the invasion of dead nutritional pathogens [12,37]. In our study, ’a.k.a. Favorita’ was inoculated with the typical necrotic pathogen (*Alternaria.solani*). Although the content of JA-Ile was significantly reduced, DEGs analysis revealed a positive reaction of JA signal pathway genes. The basic helix (bHLH) transcription factor (TF) *MYC2* occupies a high level in the JA signal transduction pathway and actively induce the transcription of downstream defense genes [27]. Although our results did not find the differential expression of *MYC2, MYC1* was down regulated at three time points. Recent studies have shown that *MYC2* activates a small amount of JA-induced bHLH protein (*MTB1, MTB2* and *MTB3*) to counteract the function of the MYC2-MED25 transcriptional activation complex, thereby negatively regulating tomato-mediated JA-mediated transcription [28,38]. Interestingly, After Aso infection, the expression patterns of *MTB1* and *MTB2* at the three time points were similar, while *MTB3* was significantly inhibited at the second time point (Supplementary Table S2). The differences in expression of the three MTB members at different time points may be due to MTB participates in the strong interaction of environmental factors, can produce a response similar to the oscillation of the biological clock during the day and night, and precisely regulate the transcription of *MYC2* and the response downstream of the JA signaling pathway. In conclusion, 6184, 10887 and 8109 DEGs were obtained at 24hpi, 36hpi and 48hpi by RNA-Seq. The functional annotation and cluster analysis of DEGs showed that in the “a.k.a. Favorita” response to Aso, the defense response mediated by JA and SA signals was activated. WGCNA analysis identified a hormone associated with co-expression modules. Our results indicate that the JA response pathway and other signaling pathways are involved in the response of ’a.k.a. Favorita’ to *Alternaria.solani*.

## Materials and Methods

### Inoculation of potato plants with *Alternaria solani*

The seed potato tubers “cv Dutch 15” (a.k.a. Favorita) were provided by Hebei Huigu Agricultural Science & Technology Co. Ltd. (Shijiazhuang, China). Individual potato tuber was sowed in pot containing a 2:1 mixture of garden soil and vermiculite without chemical fertilizer. The plant were grown under 15 hours of natural and artificial light (140 μE) for 35 days in a greenhouse with temperature adjusted to 25 °C. *A. solani* strain HWC-168 was cultured on PDA plate for mycelial growth. Then the mycelium was inoculated into tomato juice medium, and Aso sporulation was induced by exposure to ultraviolet radiation for 10 mins. Exposed to ultraviolet light for 5 days, microscopic observation confirmed spores. The spore suspension was adjusted to about 1.0×10 ^5^ spore ml^-1^, and 0.1% (V/V) Tween 80 was added to spray to the plants, and sterile water was sprayed to the leaves as a control treatment. The inoculated plants were kept in darkness at 60% relative humidity and 25 °C for 12 hours before being transferred to the previous growth condition. Potato leaves were excised from potato plants at 24, 36 and 48 hours post-inoculation (hpi) with sterile blades, and the leaves were immediately wrapped in aluminum foil and kept in liquid nitrogen.

### Measurement of chlorophyll, JA-Ile, ABA and IAA contents

The contents of total chlorophyll (a and b) and plant hormones (JA-Ile, ABA, IAA) in the potato leaves sampled from Aso-inoculated and control plants at each post-inoculation time points were measured. Leaves from three different plant individuals were selected as biological replicates. Refer to the previous method, use spectrophotometry to determined the leaf chlorophyll (a+b) content [39]. Five leaves of Sample of 20-25mg was collected by taking at least five leaves each of non infected and infected plants. The total chlorophyll of each leaf was extracted in 80% acetone and measured as mg/g of fresh weight (FW) of leaf tissue. After the liquid nitrogen was frozen, Mayobio Biomedical Technology Co., Ltd. (Shanghai, China) was commissioned to analyze JA-Ile, SA, ABA and IAA by LC-MS.

### RNA extraction and sample preparation

After inoculation with water or *A.solani*, sample at 24, 36 or 48 hpi, collect 6 independent replicates, for each sample, pool 4 leaflets from the same plant, freeze in liquid nitrogen, and transfer to -80 °C refrigerator for later use. Total RNA was isolated using a cetyltrimethyl ammonium bromide method [40]. Equal quantities of RNA from six biological replications at each stage were pooled to construct a complementary DNA (cDNA) library. Use NanoDrop2000 to detect RNA concentration and purity, and 1% agarose gel electrophoresis to detect RNA integrity. Use Agilent2100 to collect RIN value of total RNA.

### RNA sequencing and data analysis

mRNA were purified with the oligo (dT) magnetic beads. ScriptSeq RNA-Seq Library Preparation Kit (Illumina, USA) was used for RNA sequencing library preparation of each RNA sample. Libraries of all the 36 samples were sequenced by Illumina NovaSeq 6000 platform for 2×150 cycles. RNA sequencing libraries were constructed and sequenced at the Majorbio Bio-pharm Technology Co., Ltd. (Shanghai, China). The RNA-Seq datasets have been deposited at Sequence Read Archive (SRA) under the project number PRJNA574559. Adaptor and quality trimming of raw RNA-Seq datasets were performed using Fastp v0.20.0 [41]. The potato genome sequence (SolTub_3.0, GCF_000226075.1) and corresponding genome annotation were acquired from RefSeq. The trimmed pair-end reads of each sample were aligned to the reference genome using HISAT2 v2.1.0 [42]. Reads count for each protein-coding gene was calculated using HTSeq v0.11.2 [43]. Differential expression between inoculated and control plants at each time point was determined using the R package DESeq2 v1.24.0 [44] with criteria of false discovery rate (FDR) below 0.001 and absolute log2-fold change (LFC) greater than 0.58. Enrichment analyses of Gene Ontology (GO) terms and Kyoto Encyclopedia of Genes and Genomes (KEGG) pathways were performed using the R package clusterProfiler v3.12.0 [45] with criteria of FDR below 0.05. Co-expression network analysis of RNA-Seq data was performed using the R package WGCNA v1.68 [46].

### Expression analysis using quantitative real-time PCR

Gene-specific primers for qRT-PCR were designed using the Beacon Designer 7.0 program (Premier Biosoft International, California, USA) and are listed in Supplementary Table S5. Template cDNAs were synthesized using TransScript® RT/RI Enzyme Mix (TransGen, China) from 1.0 μg of total RNAs following the manufacturer’s instructions. Super EvaGreenTM qPCR Master Mix (US EVERBRIGHT®INC) was used as the labeling agent, while *Actin*7 from *Solanum tuberosum* served as the internal reference gene. These reactions were performed on an Bio-rad iQ5 Real Time PCR System, with the reaction mixture (20 μL) containing 10 μL 2 × Master Mix, 10 μmol·L^-1^ forward and reverse primers (0.4 μL each), and 1 μL template cDNA. The PCR program in this case was 95°C for 1 min, followed by 40 cycles at 95°C for 20s, 60°C for 20s and 72°C for 40s.

## Figure legends

**Supplementary Table S1**. Summary of RNA-seq datasets

**Supplementary Table S2**. Differentially expressed genes (DEGs) of potato in mock inoculated versus *Alternaria solani* (Aso) inoculated leaves at each post-inoculation time points

**Supplementary Table S3**. Results of GO term and KEGG pathway enrichment analyses

**Supplementary Table S4**. DEGs related to different physiological aspects

**Supplementary Table S5**. Primer sequences used in qRT-PCR

**Supplementary Figure S1**. Effects of spraying spore suspensions of *Alternaria solani* (Aso) isolate HWC-168 on potato foliage (Aso^+^) in greenhouse. Leaf chlorosis developed on the Aso-innoculated plants (Aso^+^) after 24 hours post-inoculation in comparison with mock-inoculated controls (Aso^-^).

**Supplementary Figure S2**. Disease symptoms of early blight in leaves of potato plants developed after *Alternaria solani* inoculation. The photos were taken at 0, 24, 36, 48 and 96 hours post-inoculation.

**Supplementary Figure S3**. Principal component analysis (PCA) of normalized RNA-Seq read counts after variance stabilizing transformation (VST) in DESeq2. Sample to sample distances are illustrated on the first two principal components (PC) comprising approximately 88% of the variation.

## Acknowledgments

This work was supported by the Earmarked Fund for Modern Agro-industry Technology Research System (CARS-09-P18), the National Key Research and Development Program of China (2018YFD0200806), and the Earmarked Fund for Modern Agro-industry Technology Research System in Hebei Province, China (HBCT2018080205).

## Author contribution

Jiehua Zhu and Zhihui Yang designed experiments. Mengjun Tian, Jinhui Wang, Chen Wang and Zheng Li performed experiments. Mengjun Tian, and Jinhui Wang analyzed data. Mengjun Tian and Jinhui Wang wrote the manuscript. All the authors read and approved the final version of the manuscript.

